# Gnotobiotic OMM^12^ mice support sustained *Clostridioides difficile* colonization without antibiotic pretreatment

**DOI:** 10.64898/2026.10.27.762127

**Authors:** Devon E. Conti, Eva Chatonnat, Quentin Lamy-Besnier, Thirishika Manivannan, Aline Crouzols, Léo C. Caulat, Céline Mulet, Marion Bérard, Laurent Debarbieux, Bruno Dupuy, Julian R. Garneau

## Abstract

Antibiotic-induced disruption of the intestinal microbiota predispose individuals to *Clostridioides difficile* infection (CDI). Although several animal models reproduce acute CDI, sustained colonization in the presence of a defined resident microbiota remains challenging to study. Here, we show that gnotobiotic OMM^12^ mice support sustained colonization for at least 20 days following oral challenge with either the reference CD630Δ*erm* strain or the clinical R20291 (RT027) strain, without prior antibiotic treatment. Early infection induced transient weight loss, rapid intestinal inflammation and transient shifts in the relative abundance of resident bacterial species. By day 20, inflammatory markers had largely resolved, and the resident microbiota had shifted back towards its baseline composition, despite continued detection of *C. difficile* vegetative cells, spores and toxins. Early bacterial community perturbations were not accompanied by detectable changes in the resident prophage induction. Moreover, whole-genome sequencing of *C. difficile* clones recovered at day 20 showed no detectable gut-specific genomic adaptation relative to the inoculum. Similar colonization kinetics and clinical outcomes were observed across independent experiments conducted over five years, supporting the consistent reproducibility of this model. Together, OMM^12^ mice provide a defined, antibiotic-free and experimentally tractable framework to investigate host-microbiota-pathogen interactions during sustained *C. difficile* colonization and a robust platform for therapeutic development.

---

*Clostridioides difficile* infection (CDI) is the leading cause of healthcare-associated infectious diarrhea and imposes a substantial clinical and economic burden worldwide ^1,2^. CDI is further complicated by recurrences, which may affect up to 45% to 65% of patients, and is associated with substantial morbidity, mortality, and healthcare costs ^3^. The primary risk factor for CDI is antibiotic-induced depletion of the gut microbiota, which reduces colonization resistance by protective gut commensal bacteria ^4–6^. The clinical manifestations of CDI range from mild diarrhea to life-threatening pseudomembranous colitis. However, asymptomatic gut colonization of *C. difficile* has also been reported in healthy adults, which may contribute to transmission and subsequent CDI in some individuals ^7^. While the adult gut microbiota restricts *C. difficile* growth and pathogenic potential, neonates are frequently asymptomatically colonized (25-70%), with rare reported cases of bloody diarrhea ^8,9^. To date, many experimental studies have focused on the acute phase of CDI, while the mechanisms involved in sustained colonization are much less known^10–12^.

Nevertheless, recurrent CDI and sustained *C. difficile* colonization have been investigated using several murine models, each with distinct experimental limitations ^13^. In germ-free mice, disease severity can range from mild symptoms to rapid death, depending on the challenge strain and experimental condition ^5,14^. In conventional mice, *C. difficile* colonization is often transient and generally requires antibiotic pre-treatment to disrupt the protective function of the gut microbiota, partially mirroring clinical risk factors. Differences in antibiotic regimens and baseline microbiota composition across mouse colonies and facilities can influence *C. difficile* susceptibility, clearance and disease severity, making direct comparisons of infection outcomes across studies difficult ^15^.

Defined microbial communities introduced in germ-free mice provide an experimentally controlled framework for investigating host–microbiota–pathogen interactions *in vivo*. The Oligo-Mouse-Microbiota (OMM^12^) is a gnotobiotic mouse model comprising 12 sequenced, mouse-derived bacterial strains spanning five major phyla represented in the murine gut microbiota ^16^. This 12-strain consortium is stably transmitted across mouse generations and can be reproducibly established in independent gnotobiotic facilities, providing a standardized microbial background for longitudinal infection studies^16,17^. OMM^12^ mice enable controlled pathogen challenge without prior antibiotic perturbation of the resident microbiota and has been used to study colonization resistance to *Salmonella enterica* serovar Typhimurium and anti-*Salmonella* phage treatment as well as diverse questions in infection biology, immunology and microbial ecology ^17–19^. Its defined composition also permits controlled addition of commensal species: *Escherichia coli* has been introduced to investigate its effects on host and community functions, whereas *Clostridium scindens* restored intestinal bile-acid 7α-dehydroxylation activity and reduced early *C. difficile* colonization and pathology ^20,21^. In addition, OMM^12^ hosts a resident prophage community whose dynamics can be studied alongside bacterial-community changes during pathogen colonization. Gut-specific viral signatures indicate that 11 prophages are stably induced by resident OMM^12^ bacteria *in vivo* ^22^. Here, we show that OMM^12^ mice support robust and sustained *C. difficile* colonization for 20 days with both the reference CD630Δ*erm* and the epidemic R20291 (RT027) isolates previously reported as low and moderate virulent strains, respectively. This model provides a reproducible and experimentally tractable system to study changes in the resident bacterial community, in pathogen genomic adaptation, host-inflammatory response and prophage dynamics during *C. difficile* sustained colonization. By avoiding antibiotic-mediated perturbation of the resident microbiota before pathogen challenge, the OMM^12^ model complements conventional murine CDI models in which colonization and disease outcomes can be strongly influenced by antibiotic and diet regimens ^23–25^.

To establish *C. difficile* colonization in OMM^12^ mice, animals were orally gavaged with 1×10^5^ spores of the reference *C. difficile* CD630Δ*erm* strain (**Figure 1A**). Colonization levels were monitored over time by quantifying vegetative cells and spores in feces (**Figure 1B**). By day 3 post-challenge, vegetative cells loads reached 10⁶ - 10^8^ CFU/g of feces, consistent with the timing of colonization establishment reported in conventional mouse models ^26^, and remained detectable through day 20 (**Figure 1B**). Spores were also detected from day 3 until day 20. Although spore shedding was more variable between animals at days 3 and 7, it became more consistent from day 10 onwards. By the day 20 endpoint, vegetative cell and spore loads were comparable in both cecal and colonic contents (**Figure 1C**). This 20-day period of sustained colonization provides an extended longitudinal window that is difficult to achieve in other *in vivo* models with an unperturbed complex microbiota, where bacterial clearance is variable and typically occurs within 6–10 days ^27^. Comparable durations of colonization in conventional models generally require microbiota perturbation through dietary manipulation, antibiotic pretreatment, or both ^23–25^. Sustained colonization of CD630Δ*erm* strain in OMM^12^ mice was reproducible across four independent experiments conducted over several years (**Figure 1B, S1A-C**).

**Figure 1.**
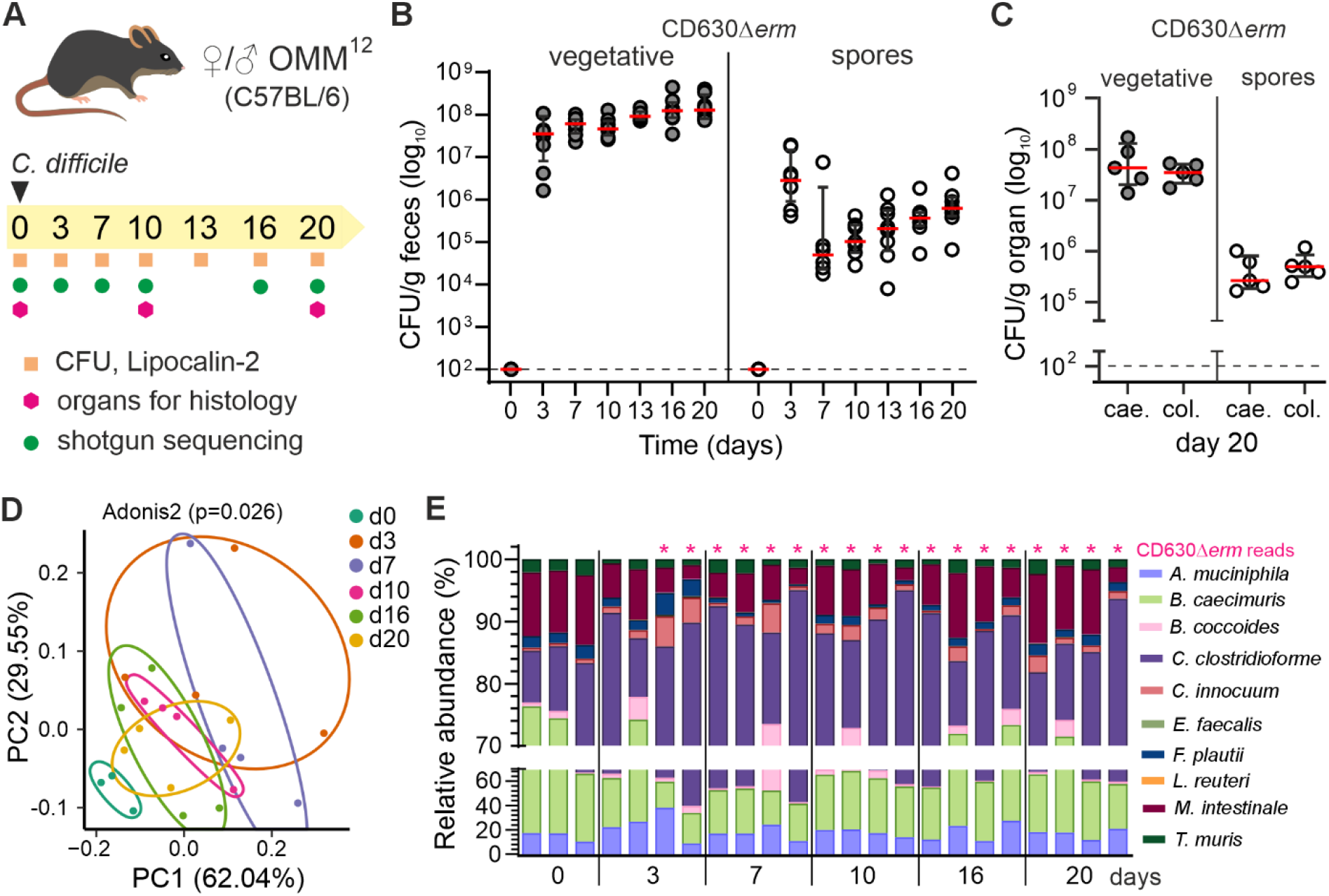
*C. difficile* establishes sustained colonization and transiently perturbs the resident bacterial community in OMM^12^ mice. **(A)** Experimental design and sampling timeline. Six- to eight-week-old male and female OMM^12^ mice were orally gavaged on day 0 with 1×10^5^ spores of *C. difficile* strains. Fecal and organ samples were collected at the indicated time points for CFU enumeration, lipocalin-2 quantification, histological analysis and shotgun metagenomic sequencing. **(B)** Quantification of *C. difficile* vegetative cells and spores in feces over 20 days following challenge with strain CD630Δ*erm*. Red horizontal bars indicate medians, and vertical error bars indicate interquartile ranges (n=8 mice). **(C)** Vegetative cells and spores count in cecal (cae) and colonic (col) contents collected at day 20 post challenge. Red horizontal bars indicate medians, and vertical error bars indicate interquartile ranges (n=5 mice). The dotted horizontal line indicates the limit of detection. Values below the limit of detection were plotted at 10^2^ CFU/g**. (D)** Principal Coordinates Analysis (PCoA) of the resident bacterial community over time, representing Bray-Curtis distances calculated from shotgun metagenomic sequencing data from feces (*C. difficile* excluded). Samples are coloured by collection day: day 0 (green), day 3 (orange), day 7 (purple), day 10 (pink), day 16 (light green), and day 20 (yellow). Ellipses were calculated using the Khachiyan algorithm. Statistical significance was calculated by Adonis2. **(E)** Relative abundances of resident bacterial species in fecal samples collected on day 0 (n=3) and on days 3, 7, 10, 16, and 20 (n=4 per time point), determined from shotgun metagenomic sequencing data using RPKM normalization and expressed as percentages. Red stars indicate the presence of reads matching the genome of strain CD630Δ*erm*. Data from panels B to D originate from an experiment performed in 2023.

To assess the contribution of the resident microbiota to this colonization profile, we compared CD630Δ*erm* challenge in OMM^12^ and axenic mice. Axenic mice maintained stable fecal vegetative cell loads until day 20, although spore loads, which were slightly more abundant at day 3 than in OMM^12^ mice, dropped progressively from day 7 onwards (**Figure S1D**). Despite sustained vegetative-cell colonization, the declining spore shedding in axenic mice indicates an unstable colonization profile. We next tested whether sustained colonization in OMM^12^ mice was restricted to CD630Δ*erm* or could also be obtained with another strain. Mice challenged with the same load of spores of the epidemic R20291 strain maintained stable fecal loads of both vegetative cells and spores through day 20, at levels comparable to those observed with CD630Δ*erm* (**Figure S1C,E**).

Together, these findings show that OMM^12^ mice reproducibly support comparable 20-day colonization profiles for two distinct *C. difficile* strains, CD630Δ*erm* (RT012, clade 1) and R20291 (RT027, clade 2), characterized by sustained fecal detection of vegetative cells and spores. Unlike axenic mice, OMM^12^ mice retain a defined resident microbiota, enabling longitudinal investigation of microbiota dynamics and host responses during sustained colonization. This model may facilitate investigation of host and microbiota factors that support prolonged *C. difficile* carriage/persistence, a clinically relevant state that can occur without overt symptoms in healthy adults and is common in infants ^8,28^. We quantified the impact of *C. difficile* colonization on the abundance of the resident bacterial community of CD630Δ*erm*-infected OMM^12^ mice by shotgun sequencing. The presence of *C. difficile* led to a significant overall change (p=0.026) in the relative abundance of the resident bacterial community during infection, with the greatest difference observed between day 3 and day 7 (**Figure 1D**). This change was marked by a decline in *Muribaculum intestinale* and *Bacteroides caecimuris*, alongside an increase in *Akkermansia muciniphila, Bacteroides coccoides*, *Clostridium clostridioforme*, and *Clostridium innocuum* (**Figure 1E**). Notably, the increase of two others species belonging to the *Clostridium* genus, *C. clostridioforme* and *C. innocuum*, suggests a metabolic interaction with *C. difficile*, which is consistent with previous observations with other clostridial species such as *Paraclostridium bifermentans*, *C. sardiniense*, *C. leptum*, and *C. hiranonis* ^5,29–31^. However, the impact of the CD630Δ*erm* strain on the bacterial community is transient and appears to be mouse dependent (**Figure 1D**). By day 10, the community composition shifts back towards their basal levels and remarkably, by day 20, the shift is almost completely restored to basal abundance in all animals, despite sustained *C. difficile* colonization.

We also investigated whether these early, transient perturbations of resident OMM^12^ bacteria during *C. difficile* infection affected resident prophage induction by sequencing the fecal virome of CD630Δ*erm*-infected OMM^12^ mice at days 1 and 3 post-infection ^22^. Compared to the uninfected controls, no additional prophages from OMM^12^ and CD630Δ*erm* strains were detected above the threshold during *C. difficile* infection, and the levels of active resident prophages (continuously induced) remained unaltered (**Figure S1F**). Altogether, *C. difficile* colonization induced a transient perturbation of the OMM^12^ bacterial community, without detectable changes in the resident prophage abundances. The subsequent return of the bacterial community towards its initial state, despite sustained *C. difficile* colonization, highlights the resilience of the resident microbiota and establishes OMM^12^ as a tractable model for future studies to dissect how resident microbiota respond to and potentially shape sustained *C. difficile* colonization.

*C. difficile* challenge induced a transient decrease in body weight between days 1 and 3 post-infection with both CD630Δ*erm* and R20291 strains (**Figure 2A**). This weight loss coincided with gastrointestinal symptoms, including semi-solid to diarrheic stools and visible agitation, which are consistent with the rapid onset of a symptomatic intestinal infection observed in conventional mice (**Table S1**). Concurrently, fecal levels of Lipocalin-2, a marker of intestinal inflammation, increased and peaked at 10⁶ pg/g at day 7, before returning to baseline from day 13 through day 20 (**Figure 2B**). Overall, body-weight loss, clinical signs, and elevated lipocalin-2 levels were transient and resolved in all animals, irrespective of the infecting *C. difficile* strain (**Figure 2A-B, Table S1**).

**Figure 2.**
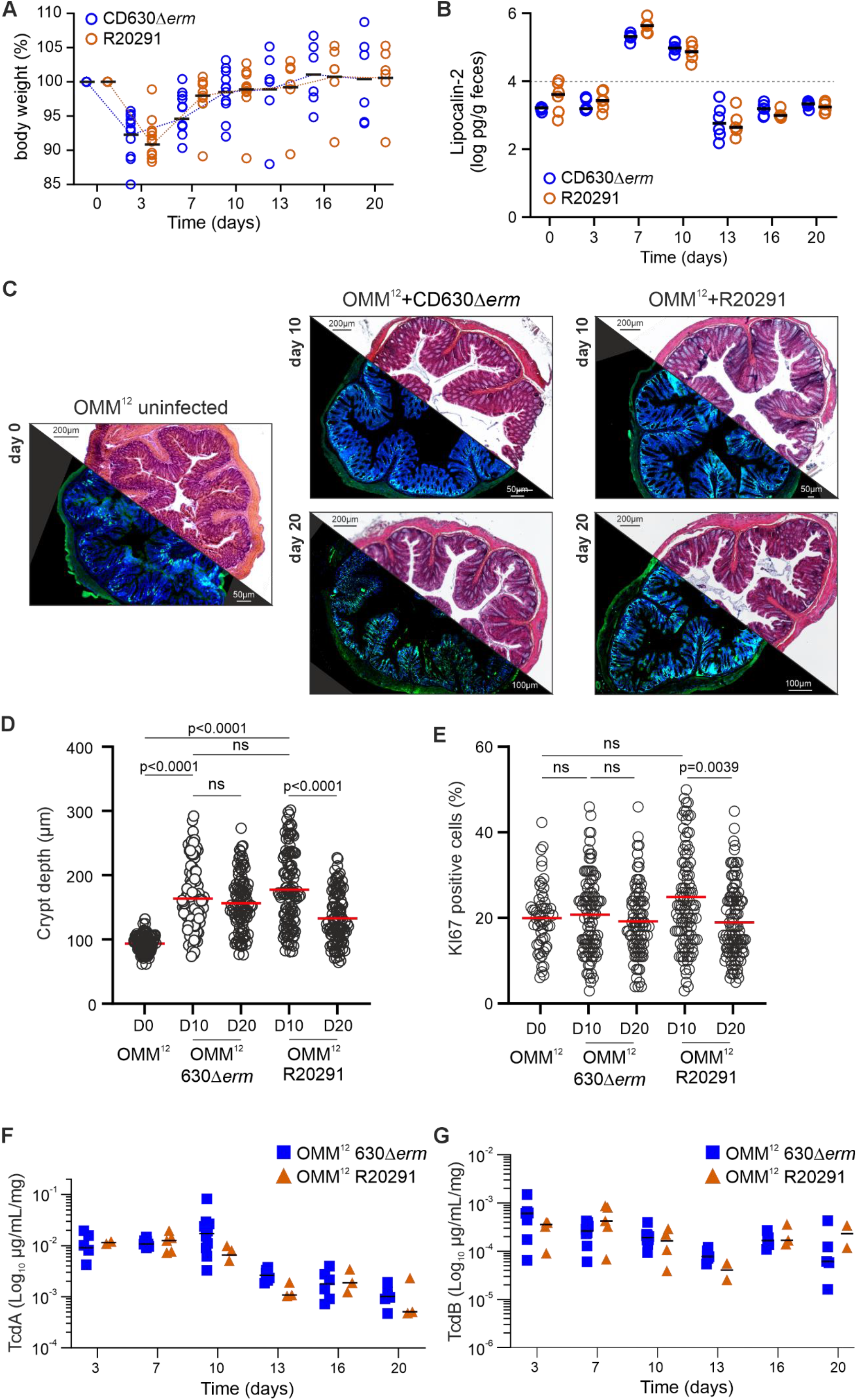
*C. difficile* challenge induces transient host responses during sustained colonization in OMM^12^ mice. **(A)** Body weight variation relative to day 0 in OMM^12^ mice infected with *C. difficile* CD630Δ*erm* (n=11) or R20291 (n=12). Black horizontal bars indicate medians. The dashed horizontal line indicates baseline body weight (100%). Some mice were euthanized on day 10 for organ collection. **(B)** Fecal lipocalin-2 levels measured over 20 days from 6 mice among those reported in panel A. Black horizontal bars indicate medians, and the dotted line indicates the inflammation threshold. **(C)** Representative hematoxylin and eosin-stained distal colon sections collected at indicated time points from OMM^12^ mice infected with the CD630Δ*erm* or R20291 strain (from panel A), with corresponding immunofluorescence images of Ki67-positive cells (green) and nuclei stained with DAPI (blue) from the same sections. Uninfected OMM^12^ mice euthanized at the start of the experiment (n=3) served as day 0 controls. Scale bars are indicated. **(D-E)** Quantification of crypt depth (D) and Ki67-positive crypt cells (E) in distal-colon sections from uninfected OMM^12^ mice and CD630Δ*erm* or R20291-infected mice collected at days 10 and 20. **(D)** Each point represents one crypt. More than 100 crypts were measured from 4 mice colon per group. **(E)** Each point represents a Ki67-positive cell. More than 55 Ki67-positive cells were quantified from 4 mice colon per group. Horizontal bars indicate medians. Statistical comparisons were performed using the Kruskal–Wallis test followed by Dunn’s multiple-comparisons correction. (**F,G**) Fecal levels of *C. difficile* toxins TcdA (**F**) and TcdB (**G**) from CD630Δ*erm* (n=5 to 10) or R20291 (n=2 to 11) infected OMM^12^ mice (from panel A) at indicated time points. Tukey’s multiple comparisons test between CD630Δ*erm* and R20291 samples at each time point did not reveal statistically significant differences neither in TcdA or TcdB levels.

Histological analysis revealed significant increased colonic crypt depth at day 10 in mice infected with either CD630Δ*erm* or R20291 compared with uninfected OMM^12^ controls, consistent with epithelial remodeling (p<0.0001 for both strains; **Figure. 2C-D**). Crypt depth decreased significantly between days 10 and 20 in R20291-infected mice (p<0.0001), but not in CD630Δ*erm*-infected mice (ns). Ki67-positive crypt-cell percentages, a marker of epithelial proliferation, did not differ from uninfected controls at day 10. Ki67 positivity declined from day 10 to day 20 in both groups of infected mice, although the decrease was significant only in R20291-infected mice (p=0.0039) **Figure. 2C-E**). While these analyses assumed that strain R20291 is more virulent than strain CD630Δ*erm*, the fecal levels of *C. difficile* virulence factors toxin A (TcdA) and toxin B (TcdB) were not statistically different (Tukey’s multiple comparisons) between these two groups of OMM^12^ infected mice, although a tendency to decrease over time was observed (**Figure 2F,G**). This contrasts with the higher level of toxins obtained *in vitro* from strain R20291 compared to strain CD630Δ*erm* (**Fig. S1G**), and supports that the host, in collaboration with its resident bacterial community, may control toxin production. Overall, these findings show that OMM^12^ mice develop an early, self-limiting symptomatic infection, characterized by transient clinical signs and intestinal inflammation, followed by sustained *C. difficile* colonization through day 20.

Finally, we assessed whether *C. difficile* undergoes detectable genomic adaptation during this prolonged colonization in OMM^12^ mice. Whole-genome sequencing performed on five pools of 10 CD630Δ*erm* clones, each recovered from an independent mouse at day 20 post-infection, detected no mutations compared to the ancestral strain. This genomic stability suggests that sustained *C. difficile* colonization in the OMM^12^ mice occurs without detectable sequence-level adaptation and is consistent with the establishment of a robust, stable host–pathogen-microbiota equilibrium.

To summarize, we found that the OMM^12^ mice address several limitations associated with the use of germ-free and conventional mice for *in vivo C. difficile* studies. In particular, the use of antibiotics or specific diet required for *C. difficile* infection in conventional mice is dispensable to recapitulate CDI in OMM^12^ mice ^23–25^. Indeed, supported by several independent experiments performed over five years, we showed that CDI initiates in OMM^12^ mice without prior antibiotic treatment and results in mild and reproducible gut inflammation followed by stable colonization for at least 20 days, unlike the unique 7-days long experiment reported with OMM^19.1^ mice ^32^. This novel animal model of CDI provides an optimal opportunity to study the mechanisms involved in *C. difficile* sustained colonization. Moreover, functional studies of the intestinal microbial community and host response characterization can be easily envisioned with OMM^12^ mice, for example, by supplementing the microbiota with beneficial bacteria to improve metabolic potential, as previously shown for other pathogens or to produce metabolites that protect against CDI ^17,20,33^. This original and robust model will also be valuable for assessing therapeutic strategies (microbial, phage-based, or pharmacological interventions) to target both initial and recurrent CDI.

---

**Figure S1.**
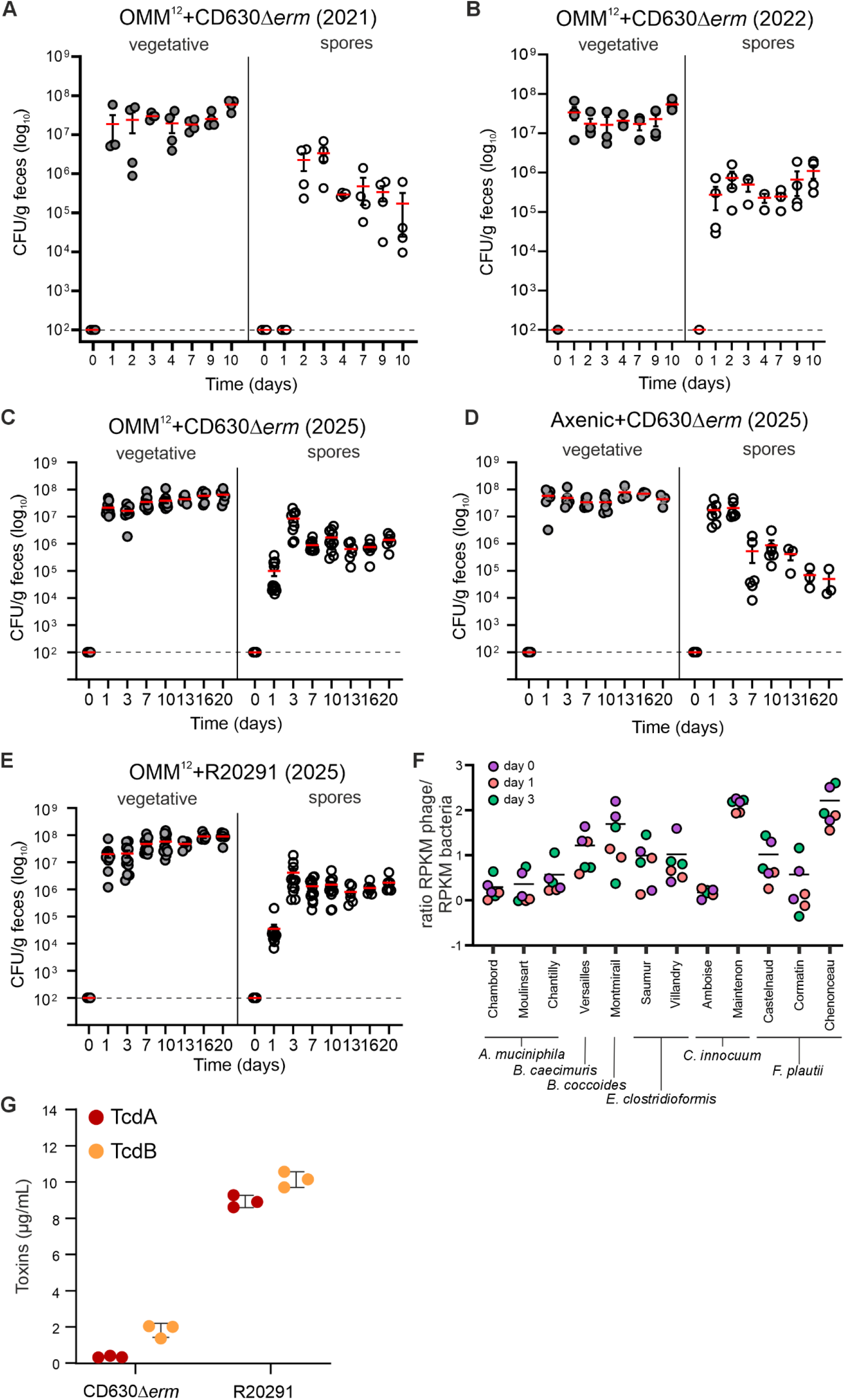
*C. difficile* colonization profile is reproducible across independent experiments and the resident prophage community is unchanged during colonization. (A–E) OMM^12^ mice were orally gavaged on day 0 with 1×10^5^ spores of *C. difficile* CD630Δ*erm* (A–D) or R20291 (E). Fecal vegetative cells (grey circles) and spores (white circles) were quantified at the indicated time points. Independent CD630Δ*erm* experiments were performed in 2021 (n=4 mice;A), 2022 (n=4;B) and 2025 (n=11; C) while R20291 experiment was performed in 2025 (n=12; E). Axenic mice were infected with CD630Δ*erm* in 2025 (n=7;D). Red horizontal bars indicate medians, and vertical error bars indicate interquartile ranges. The dotted horizontal line indicates the limit of detection. Values below the limit of detection were plotted at 10^2^ CFU/g**. (F)** Relative abundance of resident prophages in fecal viromes from CD630Δ*erm*-infected OMM^12^ mice collected on days 0 (purple), 1 (orange), and 3 (green) (n=2 per time point). Prophage abundance was calculated as the ratio of prophage abundance to host bacterial RPKM. Prophages are grouped by their predicted bacterial host, indicated below the plot. (G) *C. difficile* toxins (TcdA and TcdB) levels from the supernatant of CD630Δ*erm* and R20291 strain grown in TY broth at 37°C during 24 h.

**Table S1.** Clinical signs observed following *C. difficile* challenge in mice.

**2025** (related to Figure 2 and Figure S1C-E)
| Mouse model | Strain | N mice/sex<sup>a</sup> |  | Clinical signs and symptoms post-infection (day)<sup>b</sup> |  |  |  |  |  |  |
| --- | --- | --- | --- | --- | --- | --- | --- | --- | --- | --- |
|  |  | F | M | 0 | 3 | 7 | 10 | 13 | 16 | 20 |
| Axenic | CD630 $\Delta_{erm}$ | 4 | 3 | none | none | none | none | none | none | none |
| OMM<sup>12</sup> | CD630 $\Delta_{erm}$ | 6 | 5 | none | mo. | none | none | none | none | none |
| OMM<sup>12</sup> | R20291 | 0 | 12 | none | mo. | mi. | mi. | none | none | none |

**2023** (related to Figure 1 and Figure S1F)
| Mouse model | Strain | N mice/sex<sup>a</sup> |  | Clinical signs and symptoms post-infection (day)<sup>b</sup> |  |  |  |  |  |  |
| --- | --- | --- | --- | --- | --- | --- | --- | --- | --- | --- |
|  |  | F | M | 0 | 1 | 2 | 7 | 9 | 10 | 20 |
| OMM<sup>12</sup> | CD630 $\Delta_{erm}$ | 0 | 8 | none | none | mo. | none | none | none | none |

**2022** (related to Figure S1B)
| Mouse model | Strain | N mice/sex<sup>a</sup> |  | Clinical signs and symptoms post-infection (day)<sup>b</sup> |  |  |  |  |  |  |
| --- | --- | --- | --- | --- | --- | --- | --- | --- | --- | --- |
|  |  | F | M | 0 | 1 | 2 | 4 | 7 | 9 | 10 |
| OMM<sup>12</sup> | CD630 $\Delta_{erm}$ | 2 | 2 | none | mi. | mi. | mi. | none | none | none |

**2021** (related to Figure S1A)
| Mouse model | Strain | N mice/sex<sup>a</sup> |  | Clinical signs and symptoms post-infection (day)<sup>b</sup> |  |  |  |  |  |  |
| --- | --- | --- | --- | --- | --- | --- | --- | --- | --- | --- |
|  |  | F | M | 0 | 1 | 2 | 4 | 7 | 8 |  |
| OMM<sup>12</sup> | CD630 $\Delta_{erm}$ | 2 | 2 | none | mi. | mo. | mi. | none | none | |

<sup>a</sup> F, female; M, male.
<sup>b</sup> Animals were assessed at the indicated time points for body-weight loss, behavioral changes (including agitation), and fecal consistency (semi-solid stools, diarrhea, or blood in the stool). Clinical observations are summarized at the cohort level and reflect the proportion of animals displaying at least one clinical sign at each time point: none, no animals affected; mild (mi.), approximately 50% of animals affected; moderate (mo.), all animals affected. These categories do not indicate the severity of clinical signs in individual animals

## Methods

### Ethics statement

All animal experiments were approved by the veterinary staff of Institut Pasteur animal facility (Ref.#18.271) and the National Ethics Committee (APAFIS#26874-2020081309052574 v1). A total of 48 C57BL/6J male and female mice from 6 to 8 weeks old, bred at Institut Pasteur, were used. Animals were kept in isocages (n=2 to 4 per cage) with food and water provided *ad libitum*.

### Strains used in study and growth conditions

The *C. difficile* CD630Δ*erm* (RT012, clade 1) and R20291 (RT027, clade 2) strains were grown anaerobically (5% H2, 5% CO2, 90% N2) at 37°C in TY (3% bacto tryptose, 2% yeast extract, pH7.4) or BHI (Brain Heart Infusion, 37 g/L, Difco) medium. BHI agar plates were prepared by addition of 15 g/L of agarose. *C. difficile* vegetative cells were numerated on BHI supplemented with sheep blood (3% v/v) and antibiotics at the following concentrations: cycloserine (250 µg/mL), cefoxitin (8 µg/mL), ciprofloxacin (5 µg/mL), tetracycline (5 µg/mL). Spores were plated on CHROMID® *C. difficile* agar plates (bioMérieux) after a heat treatment (60°C, 30min).

### Spore preparation and enumeration

Spore suspensions were prepared as previously described ^34^. Briefly, 200 μL from ON cultures of *C. difficile* strains were plated on sporulation medium for *C. difficile* (SMC) medium (9% Bacto peptone, 0.5% proteose peptone, 0.15% tris base, and 0.1% ammonium sulfate) and were incubated at 37°C for 7 days under anaerobic conditions. Spores were scraped off and resuspended in 2 mL of sterile ice cold water and incubated for 7 days at 4°C. Cell fragments and spores were separated by centrifugation using a HistoDenz (Sigma-Aldrich) gradient ^35^. Spores were enumerated on BHI-Taurocholate (0.05%) agar plates and kept at 4°C on glass vials.

### *C. difficile* infection in mice

Fecal pellets from OMM^12^ or axenic mice (C57/Bl6) were collected before oral gavage with a single dose of 10^5^ *C. difficile* spores in 200 µL of PBS. Then, fecal pellets, collected at regular intervals were transferred in sterile, 1.5 mL tubes, weighed and resuspended in 1 mL of PBS. Serial 10-fold dilutions in PBS were performed and aliquots were plated onto BHI plates supplemented as reported above or CHROMID® *C. difficile* agar plates. At day 10 or 20, mice were euthanized by cervical dislocation without prior sedation and organ samples (colon and cecum) were collected. Organs were weighed and homogenized in PBS using gentle-Macs Octo Dissociator (Milteny Biotec) and serially diluted and plated as for fecal pellets.

### Fecal DNA genomic extraction

From resuspended fecal samples, 500 µL were taken and centrifuged at 8,000g for 10 min and the supernatant was removed. Pellets were diluted in 500 µL of lysis buffer (500 mM NaCl, 50 mM Tris-HCl, pH 8.0, 50 mM EDTA, 4% SDS, sodium dodecyl sulfate) and incubated for 15 min at 50°C as described previously ^36^. Then, 100 µL of lysozyme (25 µg/mL) was added and samples were incubated at 37°C for 2 h. DNA extraction was performed using the Maxwell Cell tissue kit (Promega) and samples were frozen at -20°C until use.

### Fecal virome extraction

Frozen fecal samples (2 pellets minimum) upon collection were thawed and resuspended in 14 mL Tris 10 mM, pH 8 and centrifuged for 10 min at 5,200g at 4°C. The supernatant was then filtered (0.45 μm and then 0.22 μm) and ultracentrifuged for 3 h at 270,000 x g. Pellets were resuspended in 500 μL of TN buffer (10 mM Tris, 150 M, pH 7.5). To remove free DNA and RNA, 4 U of Turbo DNaseI (Ambion, #AM2239) and 10 μL of RNAse (A/T1 mix, ThermoFisher, #EN0551) were added for 30 min at 37°C. EDTA (15 mM final), proteinase K (100 μg/mL, Eurobio) and SDS (10% final) were added and samples incubated 30 min at 37°C. The viral DNA was extracted by adding a volume of phenol chloroform-isoamyl alcohol (25:24:1) mixture, vortexing for 30 sec followed by a centrifugation for 5 min at 12,000 x g. The aqueous phase was recovered and then treated again with phenol-chloroform isoamyl alcohol similarly. Sodium acetate (300 mM final), 2 volumes of 100% EtOH and glycogen (1 μL) were added to precipitate DNA. The sample was mixed by inversion and incubated for 2 h at -80°C before centrifugation for 20 min at 15,000 x g at 4°C. The supernatants were discarded, pellets were dried and then resuspended in 20 μL Tris 10 mM, pH 8. The DNA concentration was measured with Qubit (Invitrogen).

### Library preparation and sequencing

Extracted DNA was purified using AMPure XP beads (Beckman A63882), recovered in 120 µL H2O and sonicated using Covaris (DNA 300bp) in Covaris microTUBE (Covaris, 520045). Preparation of the samples for paired-end sequencing on an Illumina NextSeq500 (2×50 bp) was performed using Invitrogen TM Collibri TM PS DNA Library Prep Kit for Illumina and following manufacturer instructions.

### Analysis of sequencing data of resident bacteria and phage community

Raw sequencing reads were first processed with fastp (v0.23.4) using default parameters to remove adapter sequences and low-quality bases, ensuring high-quality inputs for subsequent analyses. The resulting trimmed reads were then aligned to the mouse reference genome with BWA-mem2 (v2.2.1, default settings) to identify and filter out reads of mouse origin. For bacterial community profiling, Reads Per Kilobase per Million mapped reads (RPKM) was calculated for each bacterial genome by dividing the number of mapped reads by the genome length (in kilobases) and normalizing to the total number of reads per sample (in millions). The relative abundance of each bacterial species was determined as the percentage of its RPKM value relative to the total RPKM per sample, allowing for direct comparison of bacterial community composition across samples. Prophage community dynamics were similarly assessed: reads were mapped to both bacterial host genomes and their respective annotated prophage regions. RPKM values were calculated for each prophage region and corresponding bacterial host genome in every sample. The ratio of prophage RPKM to host bacterial RPKM was used to quantify the abundance or activity of each prophage relative to its host bacterium. To assess the community dynamics over time, Bray-Curtis distances were calculated using species-level relative abundance data (vegan R package). The resulting distance matrix was visualized through Principal Coordinates Analysis (PCoA), and statistical analysis was performed using PERMANOVA (Adonis2 function).

### Genomic analysis of fecal clones recovered 20 days post-infection

Glycerol stocks of fecal samples from 5 mice that had been stored at -80°C were thawed, homogenized and spread on BHI plates supplemented as described above. 10 clones isolated per mouse were collected and pooled in 100 µL of PBS. DNA was extracted using NucleoSpin® Microbial DNA kit (Machery Nagel # 740235.50) using bead beating. Thereafter, DNA was sheared by Covaris, and libraries were prepared and sequenced on an Illumina NextSeq500 (2×50 bp). Raw paired-end sequencing reads were quality-checked and trimmed using fastp (v0.23.4) using default parameters. Cleaned reads were then aligned to reference genomes CD630Δ*erm C. difficile* (LN614756.1) using Bowtie2 using paired-end (v2.5.1). Alignments were evaluated using samtools stats and samtools depth and alignment quality was summarized with MultiQC (v1.17). An in-house python script was used to visualize and confirm sufficient coverage and read depth on reference genome for each sample. To identify sequence variations between samples, the resulting cleaned reads were analyzed using Snippy (v4.6.0) for variant calling against the same reference genome. Variant reports were compiled using snippy-vcf_report to generate the HTML summary and tables containing detected SNPs and mutations.

### Lipocalin-2 assay

Frozen supernatants of fecal samples resuspended in PBS were thawed before Lipocalin-2 quantification using a commercial enzyme-linked immunosorbent assay (ELISA) kit (catalog number DY1857; R&D Systems) according to the manufacturer’s instructions.

### Histological analysis

The distal colon sample (1 cm) was fixed overnight in Carnoy (ethanol/chloroform/acetic acid 60:30:10) at 4°C for mucus preservation and then dehydrated and embedded in paraffin. All stainings were performed on dewaxed 8-μm sections by submersing them in Xylene twice for 10 min, 100% ethanol (EtOH) twice for 10 min, 90% EtOH once for 10 min, 70% EtOH once for 10 min, and PBS once for 10 min. Sections were then stained with hematoxylin and eosin. At least 20 well oriented crypts from each gut section per individual mouse were measured. The percentage of KI67 positive cells (anti-Ki67 antibodies [SP6] and goat anti-rabbit IgG H&L [Alexa Fluor® 488], both from Abcam) over total cells (DAPI) was measured on independent sections. All images were analyzed using ImageJ (Fiji).

### *C. difficile* toxins level quantification

Frozen supernatants of fecal samples resuspended in PBS were thawed on ice before ELISA assays. Wells of MaxiSorp 96-well plates (Nunc) were coated 2 hours at 4°C with 4 µg/mL of the Toxin A Rabbit Polyclonal capture antibody PCG4.1 (bio-techne) in PBS or 2 µg/mL of the Toxin B capture antibody BM347-N4A8 in PBS before washing with PBST (1xPBS + 0.05% Txeen). To block additional protein-binding sites, wells were treated for 90 min for TcdA and 60 min for TcdB with SuperBlock Blocking Buffer (Thermo Scientific) and plates were then washed with PBST. Appropriately diluted samples were distributed in the wells and incubated for 60 min at room temperature. Samples were removed and plates washed with PBST before adding polyclonal TcdA Chicken HRP-conjugated antibody LS-C128215 (LSBio) at 0.1 µg/mL in PBS or monoclonal biotin-conjugated TcdB antibody BM347-T4G1 (BBI Solutions) diluted 1:20000 in PBS and incubated 60 min at room temperature. Then, streptavidin-HRP (Thermo Scientific) at 125 µg/mL in PBS supplemented with 2% BSA was added in each well, for an additional 60 min incubation at room temperature. Plates were washed and TMB substrate solution (Thermofisher) was added in each well and kept in the dark until the development of the color (approximately 15 min). The reaction was stopped with H2SO4 (0.2 M) and the absorbance recorded at 450 nm (GloMax Discover, Promega). To quantify toxin concentrations, a calibration curve was performed with purified toxins (Sigma). For all ELISAs, the concentration of toxins was normalized by the weight of the corresponding feces. The *in vitro* quantification of TcdA and TcdB from supernatants of both CD630Δ*erm* and R20291 strains cultivated in TY broth during 24 h at 37°C was performed using the Elisa assay described above.

### Statistical analysis

The resident bacteria and prophage communities were analyzed using RPKM values from shotgun data as described above. For the bacterial PcoA analysis, Bray-Curtis distances were calculated from RPKM values (relative abundance), and the R package vegan version 2.6.4 was used for the Adonis2 multivariate analysis. Statistical analyses were performed using Prism v10.

## Data availability

The data associated to this publication will be publicly available after reviewing at https://doi.org/10.57745/BVVQ3U.

Raw sequencing reads have been deposited in the NCBI Sequence Read Archive under BioProject PRJNA1528472, comprising the fecal metagenomic data (whole-community shotgun and VLP-enriched virome libraries) and the *C. difficile* CD630Δ*erm* whole-genome sequencing data from the inoculum and from day-20 feces.

## Acknowledgements

Authors warmly thank the animal caretakers of the gnotobiology facility of Institut Pasteur and Camille Sivelle for processing shotgun sequencing. This research was funded, in part by the Groot-21 fund from the Department of Microbiology of Institut Pasteur, Agence Nationale de la Recherche, Grant ANR-24-CE15-7115 to LD and BD and from the “Integrative Biology of Emerging Infectious Diseases” (LabEX IBEID) funded in the framework of the French Government’s “Programme Investissements d’Avenir” to BD. A CC-BY public copyright license has been applied by the authors to the present document and will be applied to all subsequent versions up to the Author Accepted Manuscript arising from this submission, in accordance with the grant’s open access conditions.

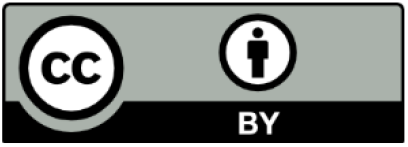

## Author contributions

JRG and QLB designed the project and secured the initial funding. AC, CM, DEC, EC, JRG, LCC, QLB and TM performed experiments and analyzed data. MB supervised the work with animals. DEC and JRG drafted the initial version of the manuscript and all the authors contributed to the final version. BD and LD secured additional funding. BD, DEC, JRG and LD supervised the project and validated the manuscript.

